## Supplementary Notes for "Inferring a spatial code of cell-cell interactions across a whole animal body"

***Specific ligand-receptor pairs drive an overall cell-cell interaction anticorrelated with spatial distance between cells.***

Since our CCI score is undirected, it can also be compared with spatial properties such as the distance between cells. Under the hypothesis that larger distances should decrease the potential of cells to interact, we expected our CCI scores to be negatively correlated with the Euclidean distances between cells. Thus, we used the distances between cells, calculated by taking the Euclidean distance between cells from a 3D atlas of *C. elegans* (Fig. N1A), as our reference data to assess our methodology and assumptions in calculating our CCI score (see ***Computing cell-cell interactions***)

We annotated each cell in the 3D atlas with a corresponding cell type in the scRNA-seq dataset (table S6), and computed the minimal Euclidean distances between each pair of cell types (Fig. N1B). The minimal distance was used because it would represent the maximal potential that cells have to interact. Thus, we next calculated the Spearman correlation between the CCI score matrix and the Euclidean distance matrix. As expected the correlation coefficient of -0.21 (P-value = 0.0016) was negative. However, the low value may be due to noise introduced by comprehensively incorporating all LR pairs into the computation, which likely includes LR pairs not necessarily encoding spatial information.

We hypothesized there is a subset of key LR pairs most relevant to spatial organization, which could be found by analyzing co-expression of cells interacting at varying proximities. To identify the LR pairs encoding spatial patterning, we ran a genetic algorithm (GA) to maximize the correlation between the CCI score matrix and the Euclidean distance matrix by randomly generating different size subsets of the LR pairs in our complete list (Fig. N2A-B). This algorithm was run 100 times, obtaining in each case a different optimal list of LR pairs due to the stochastic nature of this algorithm (Fig. N2C). Nevertheless, across all solutions, an average Spearman coefficient of -0.67 ± 0.01 was obtained (shown as an absolute value in Fig. N2B) and the maximal correlation resulted in a value of -0.70 (P-value = 1.435 x 10^-35^).

To find a core subset of LR pairs among the 100 subsets obtained by the GA, we clustered LR pairs by their co-occurrence across the GA runs and selected the cluster with members that were simultaneously present in a high fraction of the optimal subsets (Fig. N2C-D). This consensus list included 37 LR pairs (table S3), here referred to as GA-LR pairs, whose combined appearance seemed to encode proximity across cell-cell interactions, yielding a Spearman coefficient of -0.63 (P-value = 2.629 x 10^-27^) between the CCI score matrix and the Euclidean distance matrix. To test if the correlation stems from the LR pairs in the consensus list, we performed a series of permutation analyses (Fig. N3). We evaluated if the correlation computed with the consensus list was greater than the value from randomly-generated interactions between ligands and receptors in the GA-LR interactions, either by randomly permuting the ligands and the receptors (Fig. N3A) or by shuffling their labels to keep the topology of the interactions (Fig. N3B). We also subsampled the complete list of LR pairs (table S1) to obtain random subsets of similar size to the list of GA-LR pairs (Fig. N3C). In each scenario, the randomized lists yielded a smaller negative Spearman correlation than the consensus list (P-value = 0.0002, see Fig. N3).

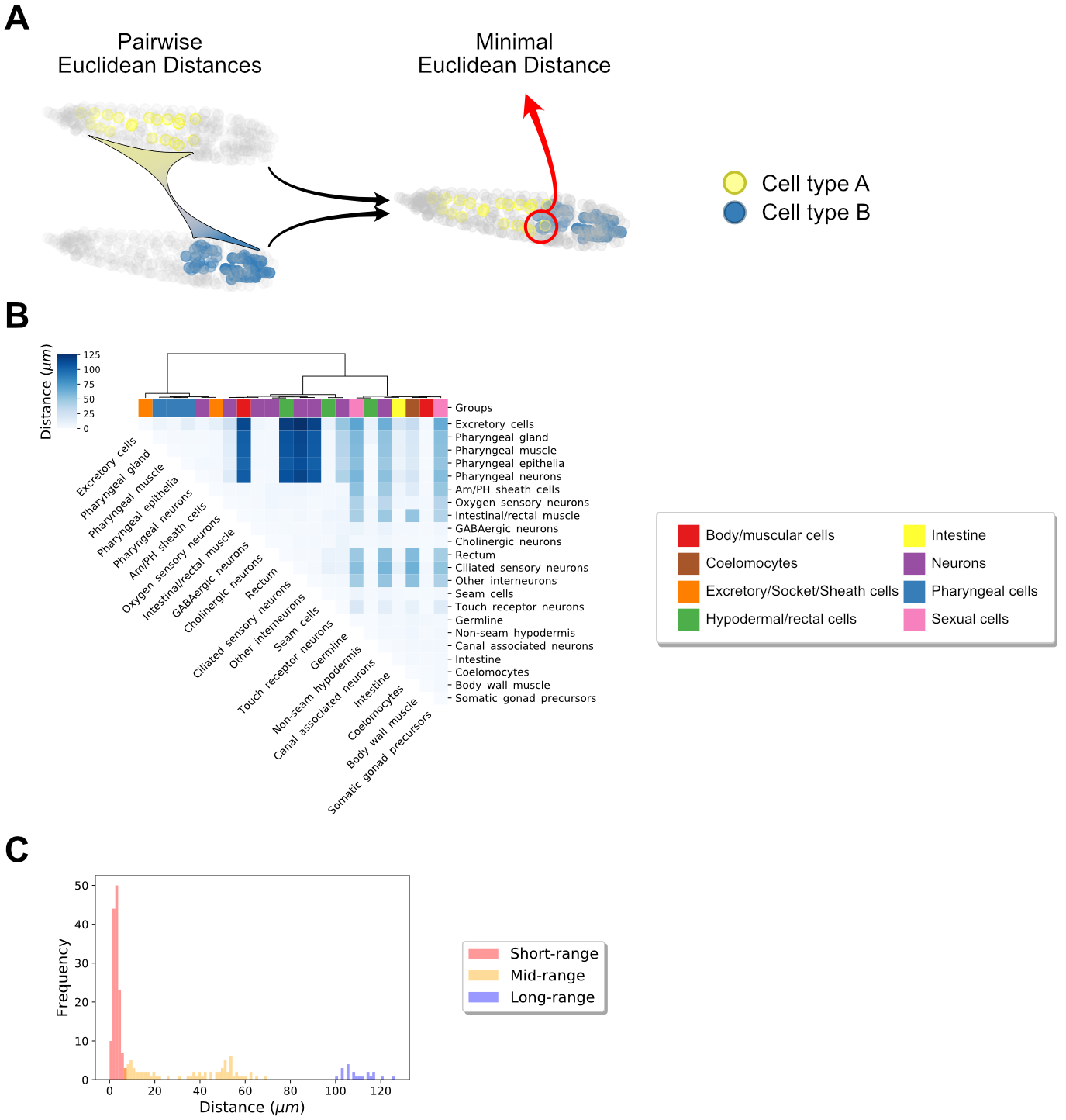

**Fig. N1. Euclidean distances between cells.**

(A) Schematic representation of computing the minimal Euclidean distance for a pair of cell types in *C. elegans*. The distance for all pairs of cells between those belonging to cell type A and those belonging to cell type B is computed; then the minimal one is selected. (B) Heatmap of resulting Euclidean distances among all pairs of cells. Depicted color key defines the eight major cell groups that were previously defined by Cao et al. (1). (C) Distribution of physical distances between cell types (built with values in the diagonal matrix in (B)). The physical distance between each pair of cell types was classified into short-, mid- and long-range distances by using a three components Gaussian mixed model.

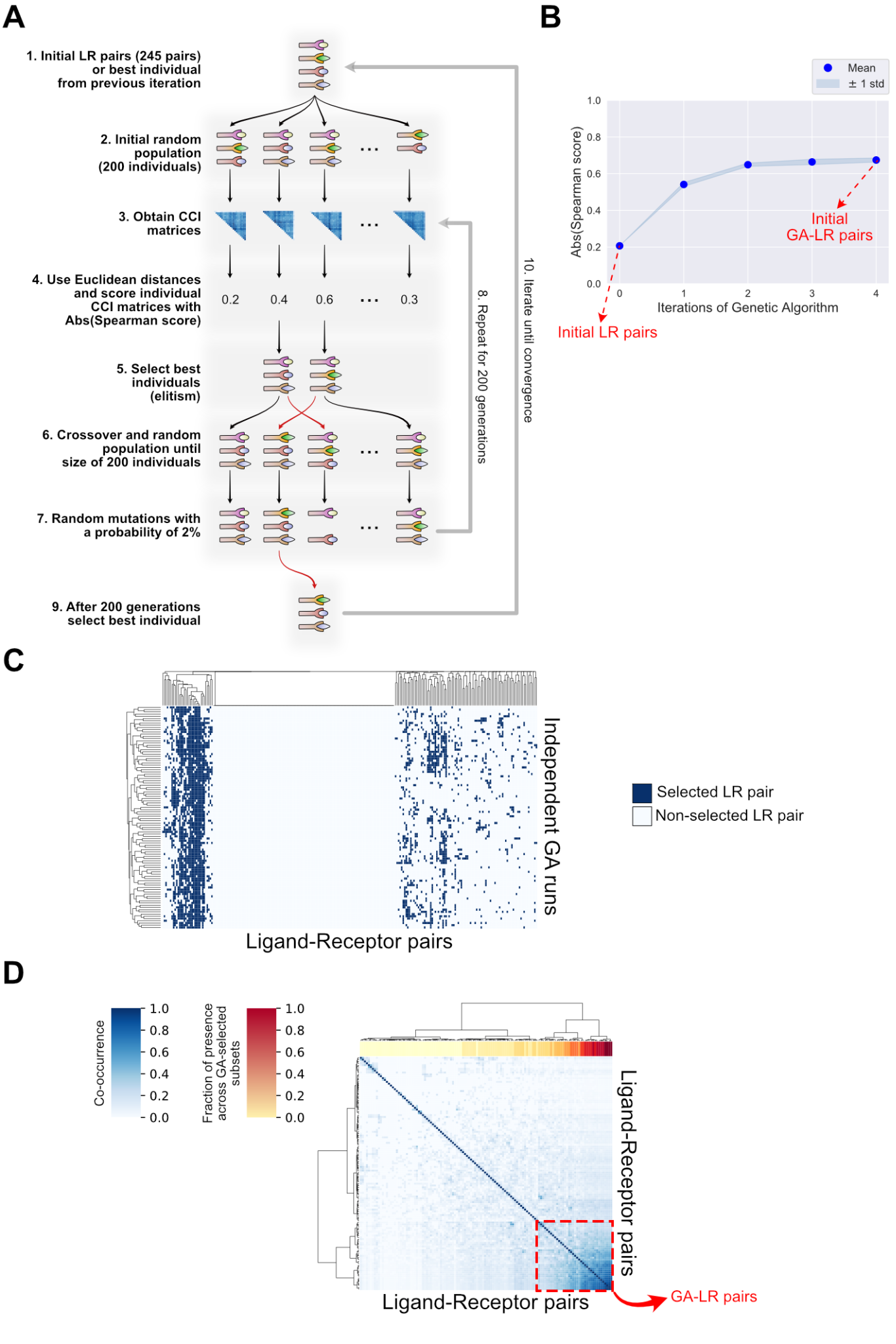

**Fig. N2. Genetic algorithm-based selection of ligand-receptor pairs leading to high CCI score-distance correlation.** (**A**) Workflow for running the genetic algorithm (GA) and selecting a subset of ligand-receptor pairs leading to an optimal correlation between CCI scores and intercellular distances. Each run consists of ten steps described in the schematic representation. This framework was run 100 times independently, leading to 100 different subsets. (**B**) Absolute value of the correlation score obtained at each iteration of the GA. 100 independent runs are plotted, showing the mean and standard deviation across them. (**C**) Heatmap indicating when each of the 245 initial LR pairs was selected by one of the 100 GA runs. (**D**) Heatmap of the co-occurrence of two LR pairs. Only LR pairs that were selected in at least one GA run are shown here. The co-occurrence is defined as the number of runs that two pairs were selected together with respect to the total number when at least one pair was selected. The dashed red square represents the cluster that was finally selected as the consensus subset of LR pairs because of the high values of co-occurrence and high fraction of presence that its interactions had across the 100 GA runs.

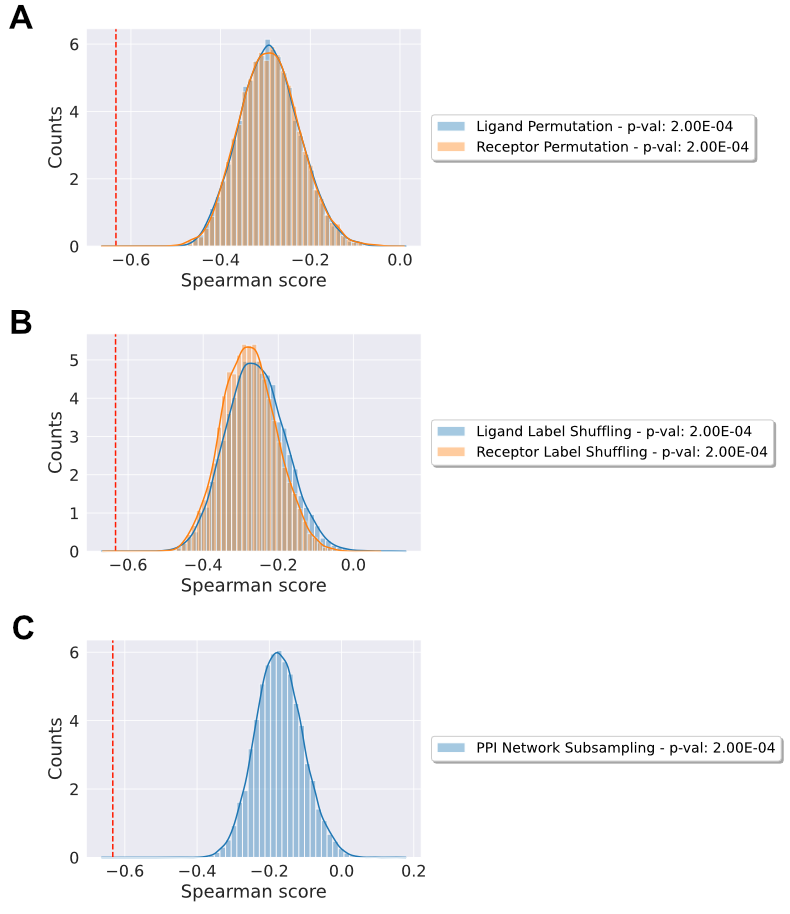

**Fig. N3. Permutation-based null distributions of the correlation obtained using the GA-LR pairs**. (**A**) Null distributions resulting from column-wise random permutation of the interactors, either ligands or receptors, separately. (**B**) Null distributions resulting from random permutation of interactor labels, either of ligands or receptors, separately. (**C**) Null distribution resulting from random subsampling of the initial LR pairs (245 interactions) to generate subsets equivalent to the GA-LR pairs (37 interactions). Each analysis in (**A-C**) was run independently 10,000 times and for each run a correlation coefficient was computed, generating a null distribution in each case. In (**A-C**) the dashed red lines represent the correlation score obtained from the consensus GA-LR pairs.

Supplementary Figures

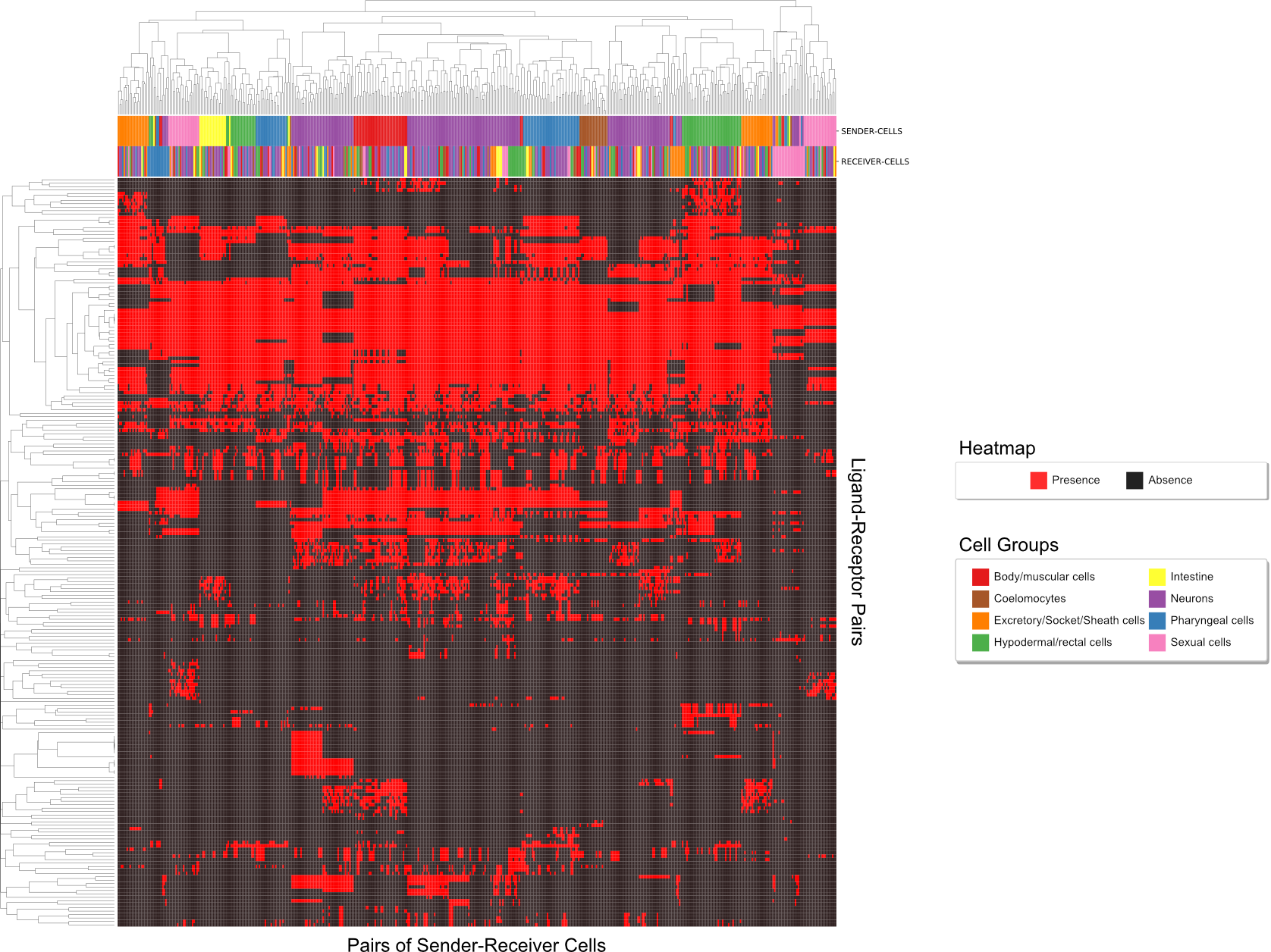

**Fig. S1. Active pairs of ligand-receptor interactions across pairs of sender-receiver cells.** Heatmap of presence or absence of ligand-receptor pairs (y-axis) across all combinations of sender-receiver cell types in *C. elegans* (x-axis). An agglomerative hierarchical clustering was performed on the Jaccard similarity for the ligand-receptor pairs (dendrogram for rows) and the pairs of cells (dendrogram for columns columns). Additionally, sender-receiver pairs were colored either by the sender cell or the receiver cell, according to the groups in the legend. Group colors were assigned previously (1).

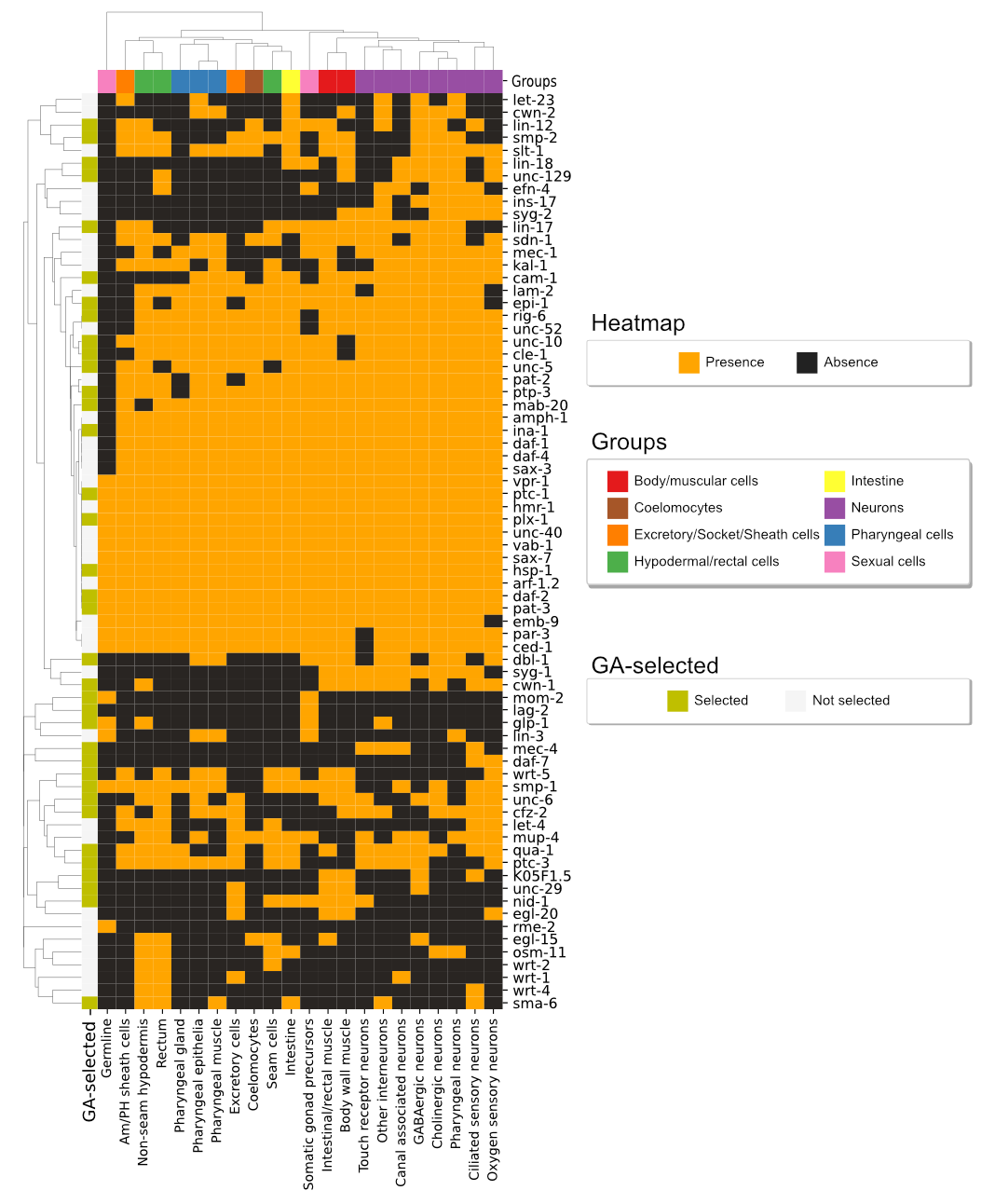

**Fig. S2. Expression of organ-phenotype associated genes in the LR pairs.** The presence or absence of proteins encoded by genes associated with organ system phenotype (y-axis) is indicated for each cell type (x-axis) according to *C. elegans* phenotype ontology. The threshold for presence is a gene expression value greater than 10 TPM; otherwise is labeled as absence. Only genes that are present in our complete list of LR pairs are shown, and members also in the GA-LR list are denoted with ochre cells (y-axis). Color keys for groups of cell types and GA-selection are depicted to the right. Agglomerative hierarchical clustering was performed using a Jaccard similarity for both genes and cell types, independently.

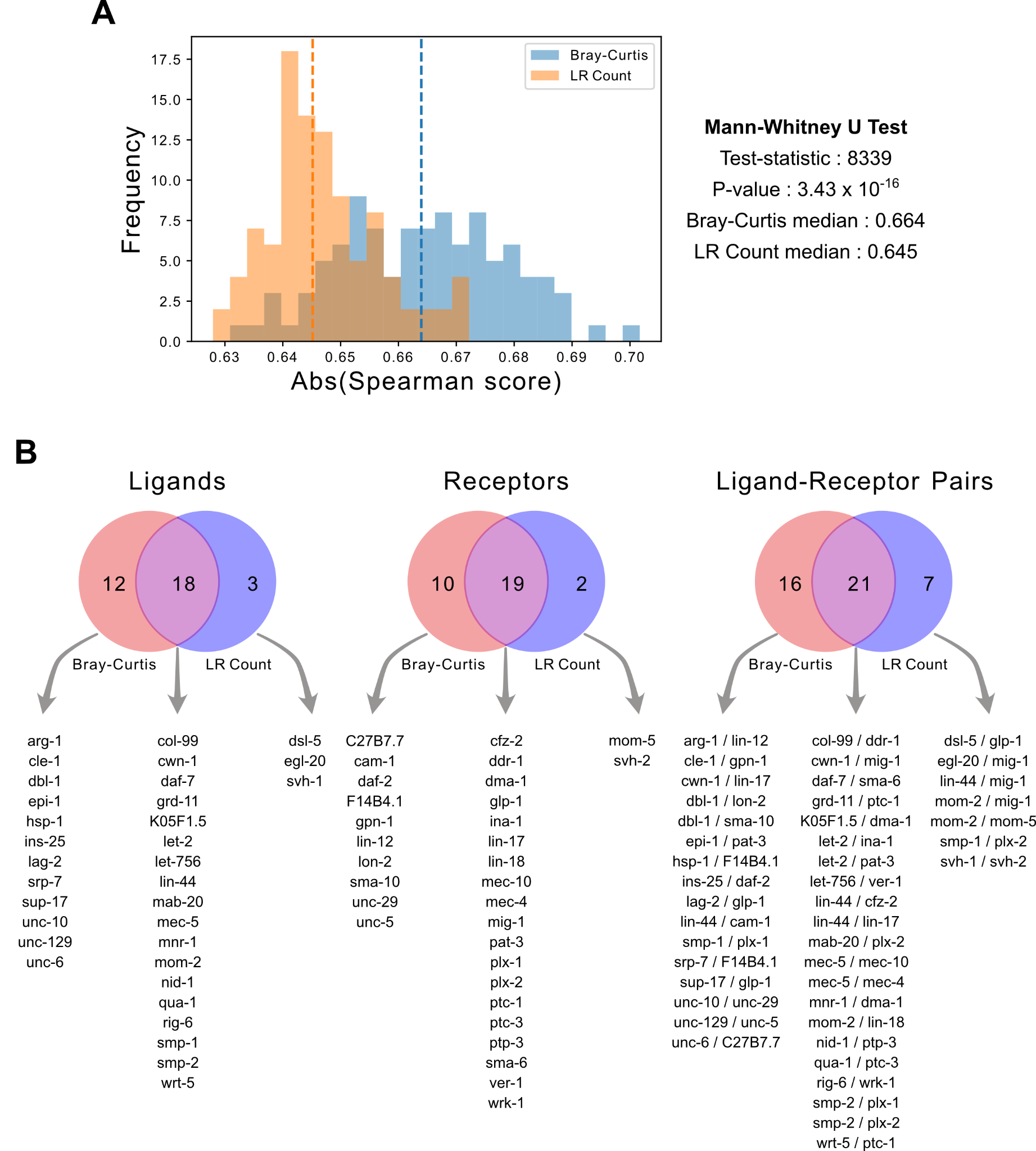

**Fig. S3. Comparison of cell-cell interaction scores used by the genetic algorithm to select ligand-receptor pairs.** Comparison of running our computational framework by using either the Bray-Curtis-like CCI score or the number of active LR pairs (LR count CCI score). (**A**) Histogram of the maximal Spearman correlation achieved in 100 separate runs of the genetic algorithm when using the Bray-Curtis and the LR count scores. The colors in the legend indicate which score each distribution corresponds to. Dashed lines represent the median values in each distribution. As indicated to the right of the histograms, a Mann-Whitney U test was performed to compare both distributions. (**B**) Venn diagrams of the ligands, receptors and LR pairs present in the consensus list of LR pairs for each of the CCI scores, obtained from the 100 separate runs of the genetic algorithm in each case. In each case, the number in the left represents the exclusive elements for the Bray-Curtis-like CCI score, the number in the right represents the exclusive elements for the LR count CCI score, and the number in the center indicates the elements that were present in both consensus lists. While lists at the bottom of the diagrams show the specific ligands, receptors and LR pairs, respectively.

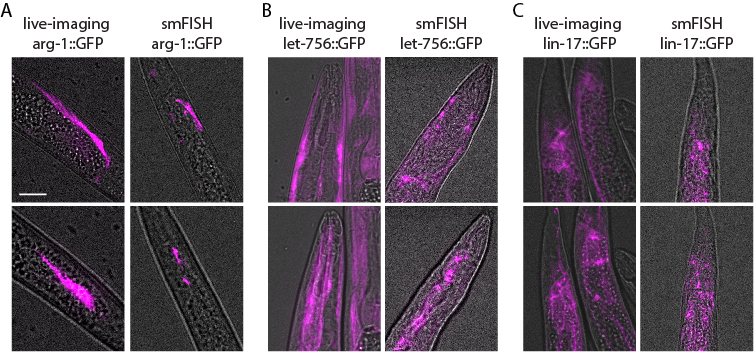

**Fig. S4. Validation of the expression patterns obtained by smFISH with GFP live imaging.** Expression patterns observed with smFISH overlap with those observed by live imaging of GFP, (**A**) *arg-1* expression in the rectal muscle in both smFISH and live imaging, (**B**) *let-756* expression in the non-seam hypodermal cells of the head in both smFISH and live imaging, (**C**) *lin-17* expression in the tail seam cells in both smFISH and live imaging. In all cases we changed the colors of the original images into magenta to make the visualizations comparable. Scale bar = 10μm.

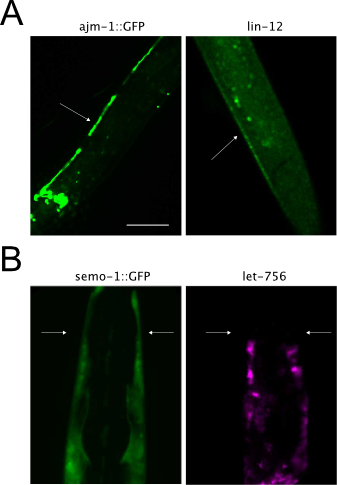

**Fig. S5. Confirmation of the localization of non-seam hypodermal cells expressing *lin-12* and *let-756.*** The expression patterns of *lin-12* in the tail (**A**) and *let-756* in the head (**B**) overlap with the expression patterns of *ajm-1* in the tail and *semo-1* in the head, confirming that the cells expressing *lin-12* and *let-756* in these regions correspond to non-seam hypodermal cells. Scale bar = 10μm.

Supplementary Files

Table S1.

Curated list of ligand-receptor interactions in *C. elegans*

Table S2.

Detailed information about ligand-receptor pairs that are used by pairs of cell types in *C. elegans*

Table S3.

Consensus list of ligand-receptor interactions selected by the genetic algorithm, corresponding to the “spatial code” of cell-cell interactions in *C. elegans*

Table S4.

Roles and experimental validation across literature of ligand-receptor pairs selected by the genetic algorithm

Table S5.

Ligand-receptor interactions of *C. elegans* described in literature

Table S6.

3D digital atlas of *C. elegans* annotated with cell types in the RNA-seq data set

**Movie S1. Tridimensional organization of cells expressing *arg-1* and *lin-12*.**

Images of the smFISH analysis projected into the 3D space. The animation shows a rotation of this projection to reflect the 3D organization of cells. Here, the intestinal/rectal muscle and the non-seam hypodermal cells expressing *arg-1* (magenta) and *lin-12* (green), respectively, are shown in the tail of *C. elegans*, as indicated in Fig. 6A.

**Movie S2. Tridimensional organization of cells expressing *let-756* and *ver-1*.**

Images of the smFISH analysis projected into the 3D space. The animation shows a rotation of this projection to reflect the 3D organization of cells. Here, the non-seam hypodermal and the amphid sheath cells expressing *let-756* (magenta) and *ver-1* (green), respectively, are shown in the head of *C. elegans*, as indicated in Fig. 6B.

**Movie S3. Tridimensional organization of cells expressing *lin-17* and *lin-44*.**

Images of the smFISH analysis projected into the 3D space. The animation shows a rotation of this projection to reflect the 3D organization of cells. Here, the seam and the non-seam hypodermal cells expressing *lin-17* (magenta) and *lin-44* (green), respectively, are shown in the tail of *C. elegans*, as indicated in Fig. 6C.
